# “An optimized pipeline for live imaging whole Arabidopsis leaves at cellular resolution”

**DOI:** 10.1101/2022.11.01.514724

**Authors:** Kate Harline, Adrienne Roeder

## Abstract

Live imaging is the gold standard for determining how cellular development gives rise to organs. However, tracking all individual cells across whole organs over large developmental time windows is extremely challenging. In this work, we provide a comparably simple method for confocal live imaging of *Arabidopsis thaliana* first leaves across early development. Our imaging method works for both wild-type leaves and the complex curved leaves of the *jaw-1D* mutant. We find that dissecting the cotyledons, affixing a coverslip above the samples and mounting samples with perfluorodecalin yields optimal imaging series for robust cellular and organ level analysis. We provide details of our complementary image processing steps in MorphGraphX software for segmenting cells, tracking the cell lineages, and measuring a suite of cellular growth properties. We also provide MorphoGraphX image processing scripts that we developed to automate analysis of segmented images and data presentation. Our imaging techniques and processing steps combine into a robust imaging pipeline. With this pipeline we are able to examine important nuances in the cellular growth and differentiation of *jaw-D* versus WT leaves that have not been demonstrated before. Our pipeline is a practical starting place for researchers new to live imaging plant leaves, but also to anyone interested in improving the throughput and reliability of their live imaging process.

## Introduction

The beautiful variety of life-forms on Earth arise from differential growth in three dimensions. Leaves offer a system to study the cellular and genetic basis of this process because they exhibit a wide range of different forms and exhibit dynamic heterogeneous growth (Avery, 1933; Poethig and Sussex, 1985; Elsner et al., 2012; Remmler and Rolland-Lagan, 2012; Derr et al., 2018; Armon et al., 2021). Advances in imaging techniques now allow us to track this development from the first few cells that initiate an organ (Calder et al., 2015; Fox et al., 2018; Kierzkowski et al., 2019; Caggiano et al., 2021). Further, cellular resolution of the same plants allows for the parameterization and fitting of models that can give greater insights into developmental processes than time-point sampling of different plants (Roeder et al., 2011; Harline et al., 2021; Strauss et al., 2022).

Yet, complex forms, like the rippling and waving leaves of the mutant *jaw-D* can stymie research by creating intractable systems for imaging (Palatnik et al., 2003). Due to its curved nature, the *jaw-D* leaf surface is particularly difficult to image in its entirety while keeping the plant alive because the leaf surface occludes itself. Similar issues arise in many other Arabidopsis mutants featuring curvature mutations, for example: *peapod, incurvata* and *curly leaf* (Goodrich et al., 1997; Serrano-Cartagena et al., 2000; White, 2006). Optical sectioning in plant tissues is often limited to the first one to two layers due to the density of plant tissue, air spaces in between cells and autofluorescence induced by chlorophyll, so imaging through curved parts is not currently feasible (Pawley, 2006). Further, even if images can be acquired, increased imaging in the z-direction comes at a time cost which can threaten sample viability. We therefore aimed to create an imaging pipeline that would minimize information lost due to tissue deformation in the z-direction while also minimizing time per sample.

In this pipeline, we have synthesized strategies from leading live and fixed imaging protocols to obtain a robust system for measuring the development of morphologically complex whole plant leaves (Grandjean et al., 2004; Reddy and Roy-Chowdhury, 2009; Hamant et al., 2019; Kierzkowski et al., 2019; Caggiano et al., 2021). Our method also makes imaging morphologically simple (relatively flat) samples easier, and permits fewer sample manipulations between imaging time points. We believe these strategies can be applied to a variety of plant tissues to improve time lapse image capture.

## Results

### Method improvements

In order to study the development of leaf primordia, we image leaves as they emerge from the shoot apical meristem. We plate seeds on phytoagar-based growth media, then allow them to germinate in the growth chamber for 2-3 days (hereafter, DAS). We then dissect the cotyledons off of the plants and allow them to recover for one day before beginning imaging (Figure 1). Before imaging we also affix a coverslip above the samples. This helps keep the samples in an ideal position for imaging. With coverslips affixed we found that perfluorodecalin is an ideal mounting media to keep samples alive and maintain image quality.

**Figure 1.**
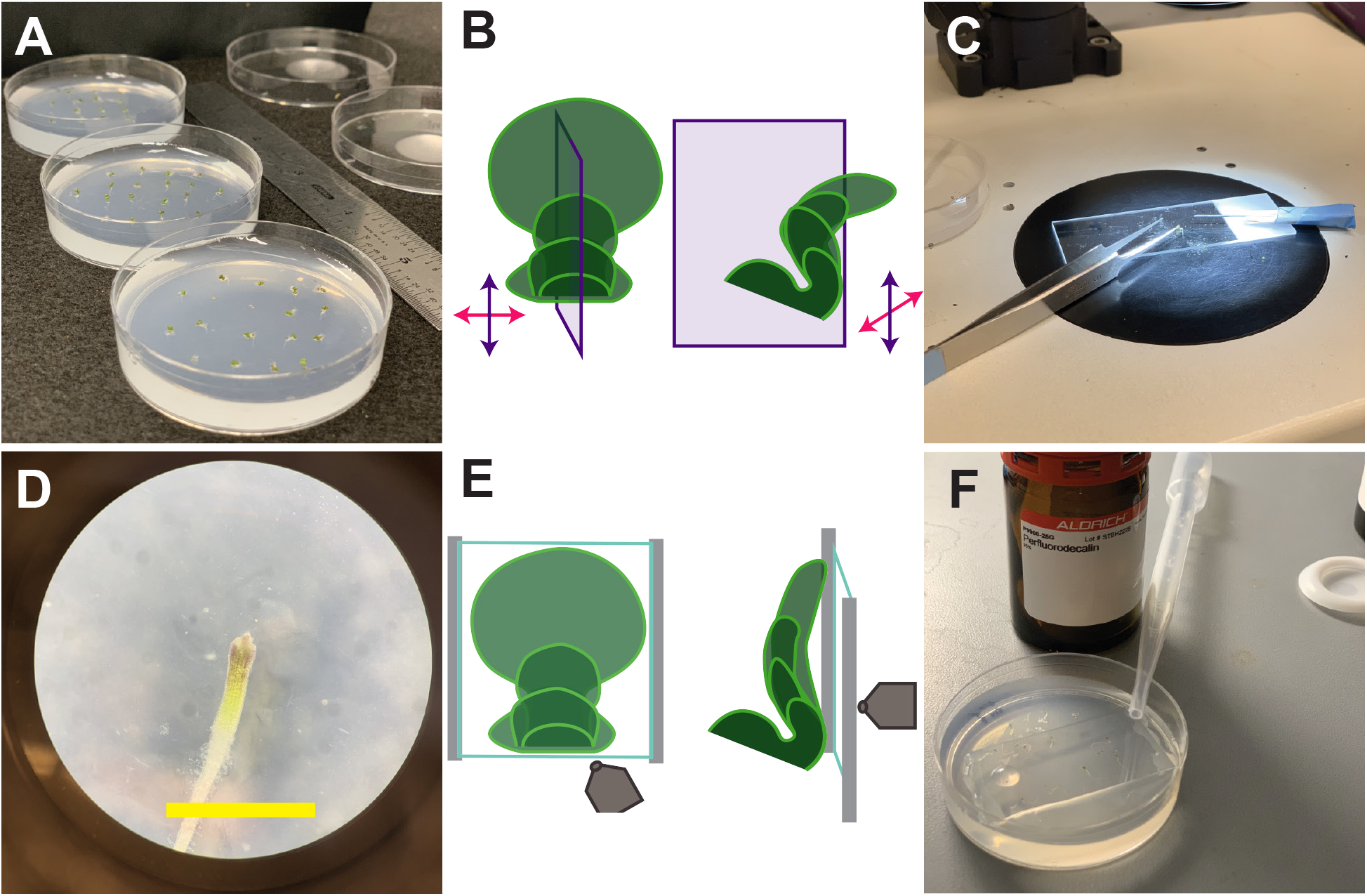
Sample preparation. (A) Germinated Arabidopsis seedlings on agar plates immediately before dissection. (B) Schematic of leaf growth without a coverslip. The abaxial surface naturally curves perpendicularly out from the plate. Directions relative to the proximal-distal and medial-lateral axes indicated with purple or magenta arrows, respectively. Purple plane indicates the cross-section displayed on the right. (C) Dissection setup under stereoscope, seedlings can be dissected within the plate or moved to a glass slide to dissect off cotyledons. (D) Stereoscope image under 50x magnification of sample post-dissection, 3 DAS (Days after sowing). Yellow scale bar = 2 mm. (E) Schematic of positions of coverslip (blue square), grease strips for coverslip suspension (gray lines) and microscope objective (gray shape, not to scale). The leaf blade flattens along the affixed coverslip. (F) One plate with all samples dissected, below suspended coverslip and immersed in perfluorodecalin (PFD).

We image the same plants this way for at least six days. Our samples contain a small plasma membrane-localized protein tagged with a fluorescent protein, so we are able to get high resolution images of every cell border in these images. This allows us to track the creation of recognizable leaf tissue containing thousands of cells from an initial unrecognizable nub of tens of cells (Harline et al. 2022). Our samples grow from hundreds of micrometers in area to millimeters in area, so they quickly exceed the single 20x imaging window at which the plasma membrane marker is resolvable (~5 DAS). We thus manually acquire tiles of smaller parts of our samples and then reassemble these individual tiles in MorphoGraphX software. We then use MorphoGraphX to convert this raw fluorescent signal into an object the computer can recognize. This involves masking the raw confocal signal, then fitting a curved surface to this mask, re-projecting the raw signal onto this surface and segmenting the signal into computer-recognized cell outlines (see Supplemental Annotated Task List). With this segmented mesh, we are able to directly measure and quantify the growth, divisions and changes in morphology of the same cell lineages throughout the imaging period. We developed scripts to speed up the processing and downstream quantification steps. The combination of these technical improvements, computational resources and our detailed supplemental information makes our pipeline ideal for researchers that are interested in tissues that curve and fold and especially newcomers to live imaging.

### Tissues grown beneath coverslips are more amenable to imaging

Plants grown in agar have a natural tendency to shift over time because the roots grow gravitropically and subsequent leaves emerge from the meristem. These developmental events shift the sample in the plate. As early leaf development proceeds, the three dimensionality of leaves becomes more apparent and imaging their entirety becomes more difficult (Figure 2). Early leaf development includes a bend that develops between the petiole and leaf blade in almost all leaves and a variety of curvature and margin patterning differences amongst mutant lines (Serrano-Cartagena et al., 2000; Palatnik et al., 2003; White, 2006; Alvarez et al., 2016) (Figure 1B, 2A, C). This can be an issue because it can lead cells from a previous time point to become obscured. These cells cannot be tracked between time points, their growth and cell division rates cannot be measured, and thus must be removed from the dataset. In order to maximize the surface of the tissue that could be imaged and tracked, while minimizing time lost to traversing z-steps, we experimented with growing plants beneath coverslips (Figure 1 D-F, Figure 2 B, D).

**Figure 2.**
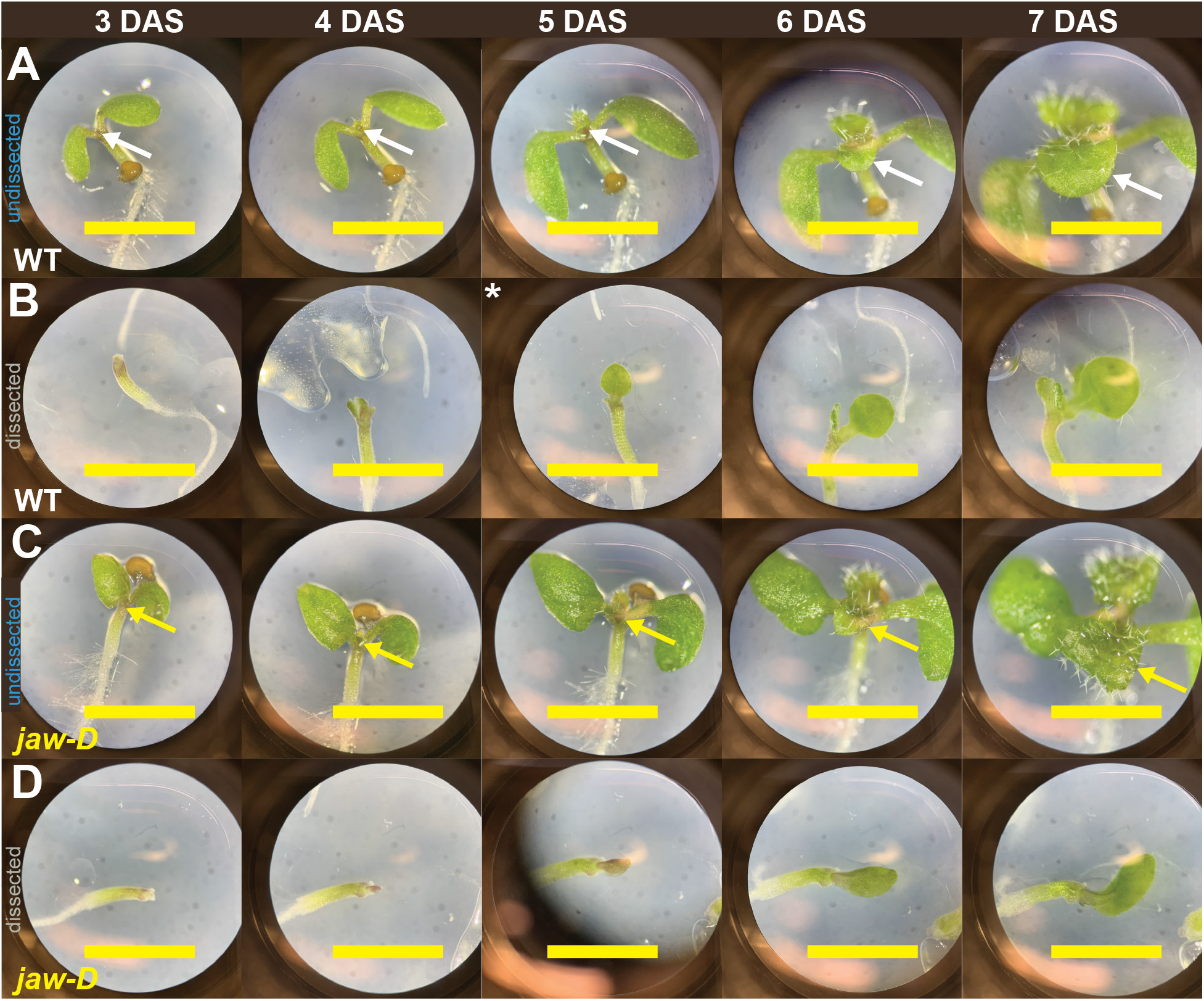
Sample growth and curving during the live imaging experiment. Stereoscope images at 50x magnification of leaf samples without coverslip or cotyledon dissection (A, C) and with dissection and affixed coverslip (B, D) for the same* WT (A-B, white) and *jaw-D* (C-D, yellow) samples over the course of a live imaging experiment. Fixed and dissected samples maintain positions with more exposed leaf tissue for imaging over time, especially in *jaw-D* where the tissue becomes curled around itself at 7 DAS. *5 DAS image was missing for this sample so an image from a different WT leaf sample is provided. Arrows indicate leaf that was imaged. Yellow scale bars = 2 mm.

Imaging and growing plants beneath coverslips offered many benefits to the pipeline. Leaves grown beneath coverslips shift much less in the plate overall and especially less in the z-dimension (Figure 2). This minimized plant movement in between imaging sessions and the risk of sample damage upon re-positioning. It also lowered the time each sample took to image by decreasing the z-step range. Further, cells were no longer lost due to tissue flipping.

Additionally, contamination of agar plates is a concern while conducting live imaging experiments. Plates will be exposed to open air for upwards of 3 hours. Some researchers use fungal inhibitors to prevent contamination (Kierzkowski et al., 2019). We have found these treatments can reduce growth. Other researchers opt to replace media regularly via complex microfluidic devices, by manual transplantation to fresh plates each day or with nutrient-minimal media (Calder et al., 2015; Fox et al., 2018; Caggiano et al., 2021). We have found that growing plants beneath coverslips radically reduces the contamination that occurs over the week or more that plants are growing. Only once in all of the week-long experiments conducted was mold found beneath the coverslip. This is a benefit as it again reduces the threat of damaging the samples from replating or losing cells by imperfect re-positioning.

### Dissecting cotyledons exposes more cells without impacting growth

Within 48 hours of being placed in the growth chamber, the cotyledons of Arabidopsis will emerge from the seed and begin to open. By this time, the first true leaves will have been initiated (Figure 2). However, due to the presence of the cotyledons, the earliest development of the first two true leaves is obscured. Previous efforts have dealt with this problem in a few ways. Regions obscured by the cotyledons have been dropped from the potential dataset (Fox et al., 2018). Images have been taken later in the development of the leaf once more of the blade has emerged (Remmler and Rolland-Lagan, 2012). Or, dissections have been performed to remove cotyledons or older leaves (Kierzkowski et al., 2019; Caggiano et al., 2021)

In accord with this last strategy, we experimented with dissecting off one and two cotyledons (Figure 1C, D, Figure 2 B, D, Figure 3). We grew WT plants with and without the cotyledons dissected in the same plates to control for condition variation (Fig 3, 4). We tested to what extent dissections improved tissue exposure for imaging and checked that growth and cell divisions were not impaired in dissected samples. Upon dissection, more cells along the early primordial margin and base are revealed and amenable to segmentation (Figure 3, 4A-D). Importantly, there is no significant difference in the areal growth or cell divisions between dissected and undissected samples (Figure 4E-H).

**Figure 3.**
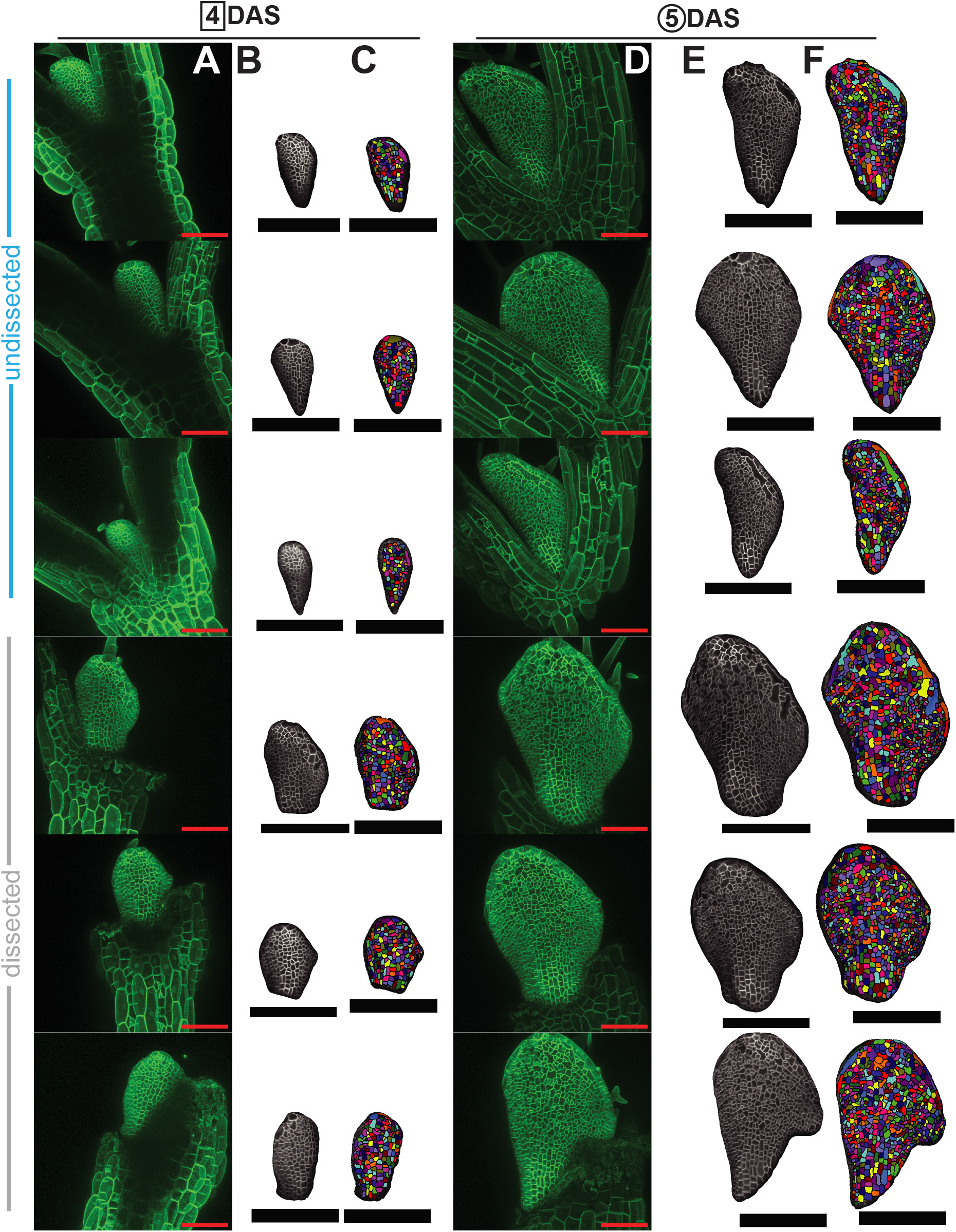
Dissecting cotyledons increases imaging accessibility. Comparison of exposed and segmentable cells in undissected (top three rows) versus dissected (bottom three rows) samples for the sample replicates from 4 DAS (A-C) to 5 DAS (D-F). More cells and more of the basal petiole and margin regions are accessible in the dissected samples. (A, D) Raw confocal images as maximum intensity projections. (B, E) Snapshots of the same images rendered as 2.5D meshes in MorphoGraphX 2.0. (C, F) Snapshots of the same meshes with segmentable cells indicated with unique colored labels and outlined in black. Red scale bars = 100um. Black scale bars = 200 um.

**Figure 4.**
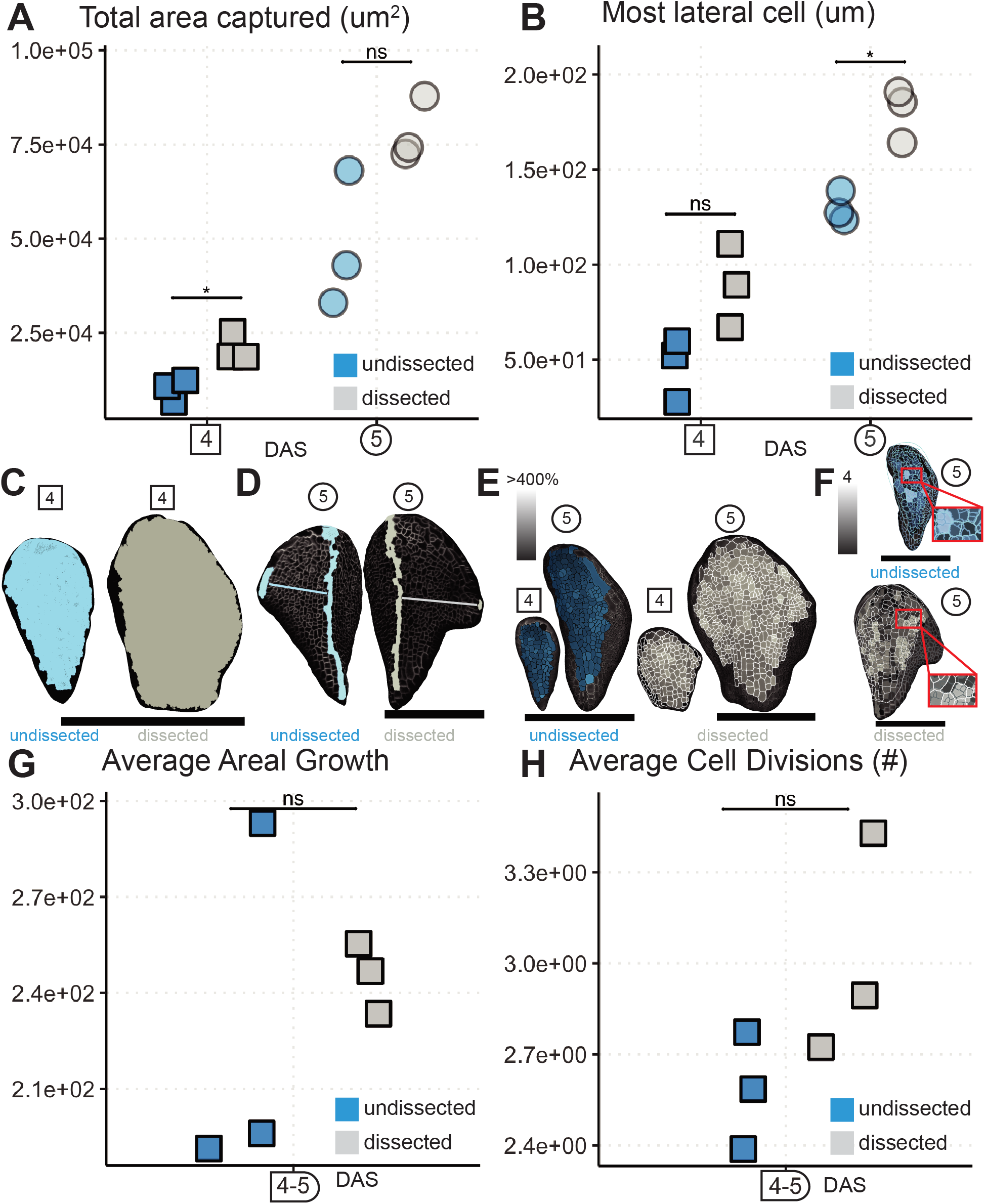
Dissecting cotyledons increases imaging accessibility without altering growth and cell divisions. Quantification of the segmentable area (A) and most lateral cell captured (B) from the three replicates in Figure 3 at 4 DAS (squares) and 5 DAS (circles). Dissected samples (gray) trend towards or have significantly more segmentable area and furthest lateral cells exposed for imaging than undissected samples (blue). (paired Student’s t-tests, * = p < 0.05). (C) The largest total area captured for each condition at 4 DAS represented on its respective mesh. (D) The furthest lateral cell for each condition, undissected (blue) or dissected (gray), at 5 DAS shown on its respective mesh. Medial cells are also selected with a line drawn for distance reference. Note how the dissected sample’s most lateral cell is lower in the tissue so more marginal cells can be captured through dissection. (E) Cell areal growth from 4-5 DAS represented on the 4 and 5 DAS meshes for the sample with the median value of average growth for each condition. (F) Cell divisions from 4-5 DAS shown on the 5 DAS mesh for the sample with the median value of average divisions for each condition. The corresponding 4 DAS mesh is scaled and overlaid to show the parent cell outline for daughter cell clones. Insets are zoomed views. (E-F) Lighter shading indicates more growth (percent area increase) or divisions (#) as indicated. Average growth (G) or number of cell divisions (H) from 4-5 DAS for each replicate. Average growth and divisions are not statistically different between treatments, so dissection does not interfere with regular development (Student’s t-test p > 0.05). Black scale bars = 200um.

### Perfluorodecalin maintains samples over many days

In our early experiments we used water based solutions to immerse the samples. The leaf growth stalled, possibly because these solutions often absorb into the media and can form a vacuum with the coverslip (Figure 5, Video 3). This prompted us to search for other immersion solutions with high refractive index to maintain good imaging resolution while not leading to tissue stalling. We attempted imaging with glycerol, iodixanol and perfluorodecalin (PFD). We found that PFD had the best results in maintaining image quality. PFD is known to permit the dissolution of gasses like oxygen and carbon dioxide, which likely contributes to its prevention of tissue stalling (Littlejohn et al., 2010). Notably, PFD is also slippery and absorbs into the media much less, so it is easier to remove in between imaging sessions.

**Figure 5.**
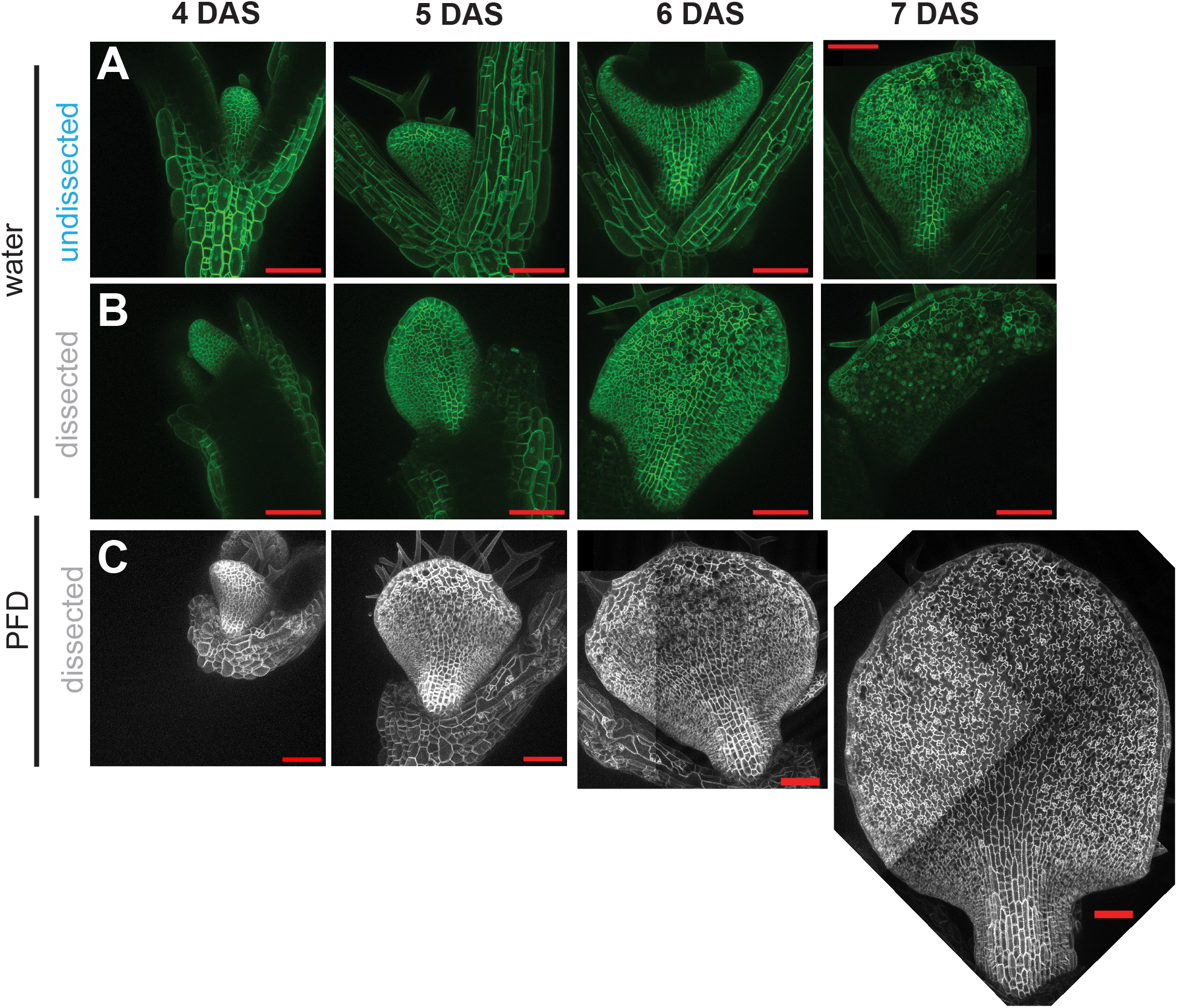
PFD allows long term imaging of samples under coverslips. Confocal maximum intensity projections of samples undissected mounted with water (A) or dissected mounted with water (B) or dissected and mounted with PFD (C) all with coverslips affixed. Samples continued to grow from 4-7 DAS only with PFD as the mounting solution. Red scale bars = 100um.

### Development of MorphoGraphX scripts increases image processing and analysis efficiency

Image processing, cell segmentation and lineage tracking can be a laborious and time intensive process (see Image Processing in the Materials and Methods and the task list in Supplemental Information for details). We therefore aimed to make the final image data analysis as efficient as possible. We used the existing MorphoGraphX infrastructure to invoke custom analysis scripts (Strauss et al., 2022). We developed scripts to call different types of mesh measurement and display processes. Our main script (iterative_growth_and_measures.py; Supplemental Code 1; https://github.com/kateharline/roeder_lab_projects/tree/master/mgx_scripts) allows users to designate which cell characteristics to measure and to specify the display and production of heatmaps with MorphoGraphX processes (Video 4). The script features pauses for measures that require user input, like selecting cells for distance measures, as well as for mesh arrangement before mesh snapshotting (Video 4).

We also developed a script (multi_resize.py; Supplemental Code 1; https://github.com/katehar-line/roeder_lab_projects/tree/master/mgx_scripts) to address an issue with files exported from ImageJ (Video 5). Sometimes the file headers are written in a way that MorphoGraphX cannot read the step size. So instead of loading an image volume, it appears as a one dimensional plane. The script iteratively opens any folder containing image files and resets the stack x,y,z dimensions to properly represent the volume, then saves the adjusted stack file. This is helpful especially if these exporting issues arise in the middle of a long term experiment when stacks need to be assembled every day to check that the entire sample was captured.

### The new pipeline enables the direct quantification of cellular mechanisms of development

Combining our imaging techniques with our custom MorphoGraphX scripts enables us to capture a large dataset encompassing the early development of WT and *jaw-D* leaves (Figure 6). From this dataset, we could analyze cell growth, division and morphology characteristics between different tissue regions, like the petiole and the margin (Figure 7 and Harline et al. 2022).

**Figure 6.**
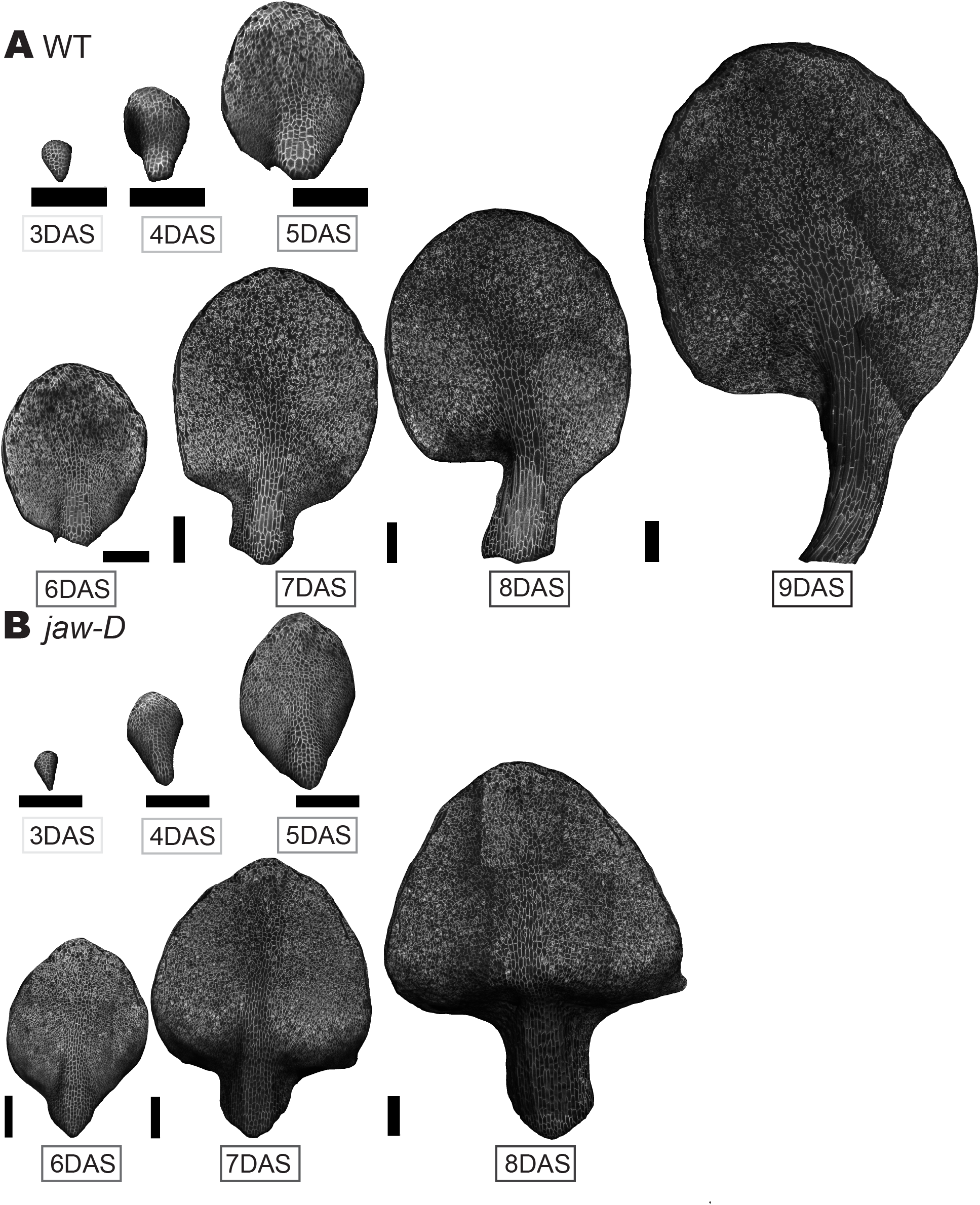
Successful imaging of complex leaf development. MorphoGraphX 2.5D mesh representations of the same WT leaf imaged from 3-9 DAS (A) and the same *jaw-D* leaf imaged from 3-8 DAS (B). The majority of the entire organ of both WT and *jaw-D* samples is visible allowing rich analysis of growth, divisions and cell types across the tissue. Black scale bar = 200um. This dataset was further analyzed in (Harline et al. 2022).

**Figure 7.**
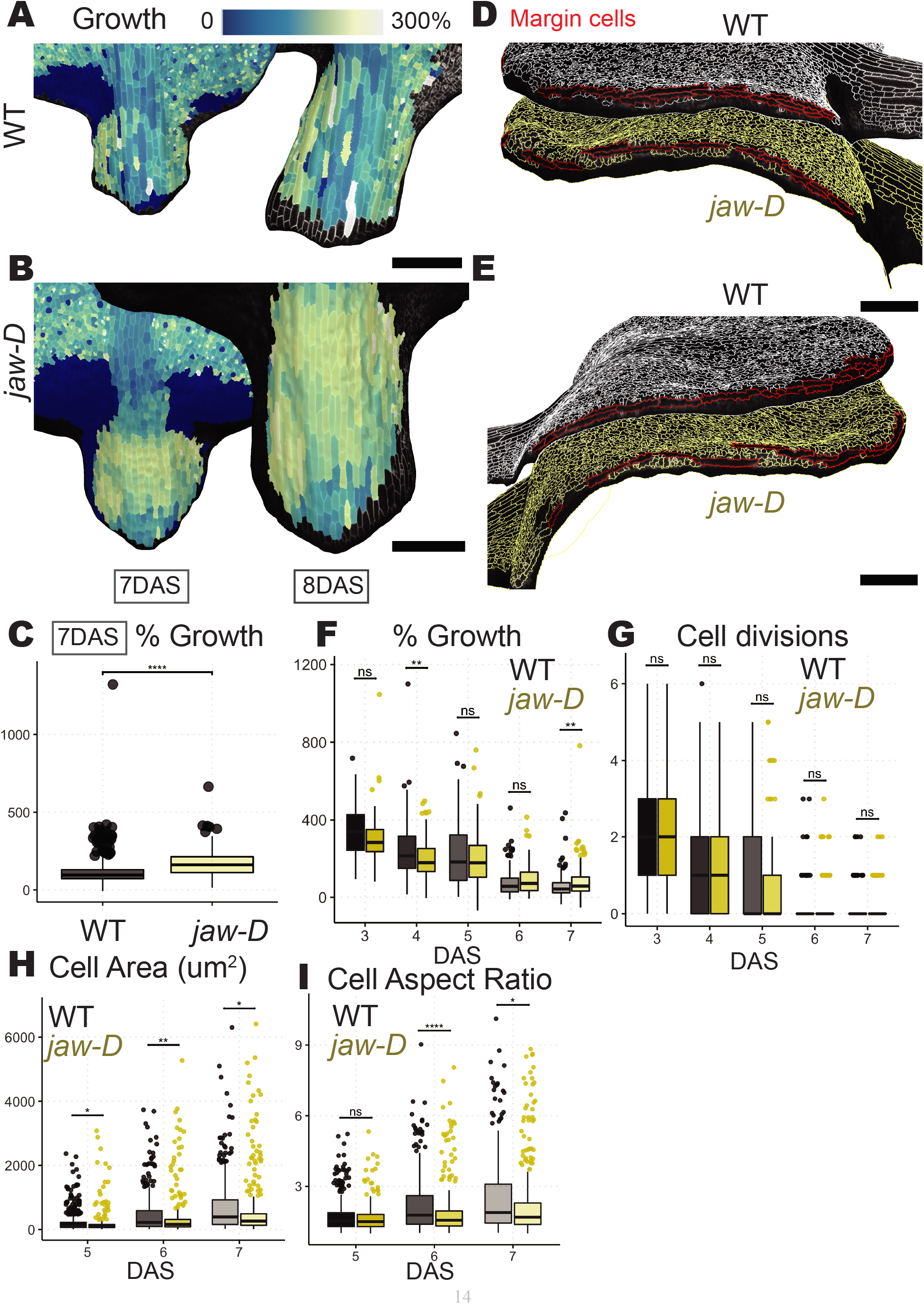
Pipeline can quantify petiole growth and margin patterning disruption in *jaw-D*. (A-B) Cell areal growth rates for 7-8 DAS displayed on 7 and 8 DAS meshes for WT (A) or *jaw-D* (B) leaf samples. (C) Quantification of areal growth rates of petiole cells. *Jaw-d* petioles have higher growth rates on average, with less variability (Student’s t-test p < .0001. CV asymptotic test p < 1.578488e-19). (D-E) Side views of WT (top, white outlines) and *jaw-D* (bottom, yellow outlines) leaves with margin cells selected in red. Elongated margin cells form a continuous border around the edge of WT leaves, but *jaw-D* leaves have gaps. (F-I) Quantification of areal growth (F), cell divisions (G), cell area (H) and cell aspect ratio (I) for all cells 10 μm from margin from 3-8 DAS (F-G) or cells 25um from the margin from 5-7 DAS (H-I). Growth and divisions are largely indistinguishable between cells at the margin between WT and *jaw-D*. Cells near the WT margin are larger and more elongated than *jaw-D*, so margin differentiation may be disrupted. (Student’s t-test * = p < .05, ** = p < .01, *** = p < .001, **** = p < .0001).

### Jaw-D *petioles exhibit more homogeneous growth*

By labeling the cells of the petiole through the script, we were able to compare the average growth rates and variability of growth in these cells. This uncovered that, at 7 DAS, the cells in *jaw-D* leaves exhibit greater average areal growth amongst cells and less variability between cells than WT (Figure 7A-C, Student’s t-test p < .0001, CV asymptotic test p < 1.578488e-19). In other work, we have shown that *jaw-D* petioles are shorter than WT (Harline et al. 2022). Our method suggests that this final organ level phenotype could be created by homogeneity and misregulation of growth patterns in the *jaw-D* petiole.

### *The* jaw-D *margin is disrupted*

We also used our cell labeling and quantification pipeline to explore the growth and morphology of cells at the leaf margin. The *jaw-D* leaf curling phenotype has been attributed to overproliferation of cells at the margin (Avery, 1933; Alvarez et al., 2016; Rebocho et al., 2017; Challa et al., 2019). Using the script, we selected a band of cells along the edge of leaves that we defined as the margin. Then, we quantified the growth, divisions, and characteristics of cells a prescribed distance away from this designated leaf edge. When we consider cells 10 μm or less away from the edge over time (the average width of cells from 3-7 DAS), we see that the average growth rates and divisions between WT and *jaw-D* leaves generally are no different (Figure 7 F-G). Only from 4-5 DAS and 7-8 DAS is the average areal growth different, and at 4-5 DAS it is actually higher in WT than in *jaw-D* (Student’s t-test p < .01).

Previously, cell cycle markers and cell density were used as a proxy for proliferation (Efroni et al., 2008; Alvarez et al., 2016; Beltramino et al., 2018). However, our direct measurement of cell divisions and morphology in leaf 1 suggests that, it may appear that there is more proliferation at the *jaw-D* leaf edge because margin cells are less well defined (Figure 7 D-E, H-I). In WT leaves the margin consists of elongated cells in a continuous band around the edge that may be stacked in multiple rows (Figure 7 D-E, top). While, in *jaw-D* the leaf edge exhibits some elongated cells, they can be discontinuous with gaps of small cells and usually are only one layer thick (Figure 7 D-E, bottom). When we quantify the morphology of cells 25um from the leaf edge (the average width of cells from 5-7 DAS), we find that WT cells are generally larger and longer on average (Figure 7 H-I, Student’s t-test * = p <.05, ** = p < .01). These results suggest live imaging and computational analysis is required to confirm the cellular dynamics that give rise to tissue morphology.

## Discussion

We provide an optimized method for capturing the relationship between cell and tissue morphology changes over multi-day time scales. We have conducted our experiments in the relatively fragile and morphologically dynamic early leaves of Arabidopsis WT and *jaw-D* mutant. Through our pipeline, we are able to characterize and quantify the entire leaf organ development at the cellular level. We demonstrate an analysis of two distinct leaf tissue regions, the petiole and the margin. This analysis suggests that growth homogeneity in the petiole and disrupted margin cell differentiation may contribute to the *jaw-D* leaf rippling phenotype. Our work emphasizes the importance and feasibility of measuring cell divisions, growth and morphology directly in living tissues to validate and discover mechanisms of development.

Our live imaging pipeline is able to capture morphologically complex tissue in a relatively straightforward and quick way. We believe that our imaging technique, processing details and scripts could be applied to a variety of systems that feature morphological complexity.

## Methods

### Plant material

WT plants are ecotype Col-0. *jaw-D* leaves are the *jaw-1D* allele from the Arabidopsis Biological Resource Center (ABRC stock number CS6948; (Palatnik et al., 2003). Plants were crossed with the epidermal specific fluorescent reporters for plasma membrane (pAR169 *AtML1:mCitrine-RCI2a*) and nucleus (pAR229 *AtML1:H2B-TFP*) (Roeder et al., 2010; Robinson et al., 2018). In subsequent generations, plants homozygous for both markers were selected. Note, only the plasma membrane marker was analyzed for the purposes of this paper.

### Growth conditions

Plants were grown in growth chambers at 22°C under continuous ~100 μmol m^−2^ s^−1^ light. Seeds were sterilized by first washing in a 70% ethanol solution supplemented with .01% SDS for 7-10 minutes on a nutating shaker, then at least three washes with 100% EtOH, then drying on sterile filter paper. Seeds were then plated on 50 mm petri plates with sterilized toothpicks. Growth media was 0.5x Murashige and Skoog media (pH 5.7, 0.5g/L MES, 1% phytoagar) supplemented with 1% sucrose. Plates were sealed with micropore tape. Plants were stratified at 4°C for 2-7 days before being placed in the growth chamber.

### *Sample preparation* (Figure 1)

Plants were harvested for dissection 2-3 days after being placed in the growth chamber (DAS = days after sowing) (Figure 1A). 0-2 cotyledons were dissected off using a BD 23g 1 and ¼ inch needle and no. 5 forceps (Figure 1C, D). One cotyledon was held with the forceps, while the needle was nestled along the adaxial side of the free cotyledon until that cotyledon was sliced off. The second cotyledon was removed in a similar manner, but with the stem gently steadied between the forceps. Plants were allowed to recover for 24 hours before imaging commenced. So imaging commenced either at 3 or 4 DAS. Before imaging, seedlings were arranged to expose the entire abaxial surface of one of the leaves. For researchers interested in comparing the first two leaves, or vegetative meristem, the sample can be placed on its side to reveal these areas. 50×22mm coverslips were then adjusted in size (strategically broken) to fit over the arranged seedlings (Figure 1F). Vacuum grease was extruded from a syringe without a needle onto both 22mm coverslip ends and then used to suspend the coverslip above the samples (Figure 1E, F). The gap between the media and coverslip was filled with perfluorodecalin (found to be effective) or water (found not to be effective), then samples were imaged (Figure 1F). In between daily imaging sessions, imaging solution was drained out from beneath the coverslip (only effective for PFD), plates were then re-sealed with micropore tape and returned to the growth chamber.

### Confocal Imaging

Plants were imaged on a Zeiss 710 Confocal laser scanning microscope with a 20x Plan-Apochromat NA 1.0 water immersion lens. The mCitrine plasma membrane marker was excited with a 514 nm argon laser and emission spectra collected from 518-629 (for the experiment in Figure 3 and 5A-B) or 519-650 nm (for the experiment in Figure 5C-7), through a 458/514/594 (for the experiments in Figures 3, 5A-B) or 458/514 dichroic mirror at 1-2% detector gain (for the experiment in Figures 5C-7). If the plants could no longer be captured within one stack, the entire visible surface of the leaf was tiled over manually using cellular landmarks adjusting the z-range to minimize time imaging the leaf. When tiled manually, stacks were assembled in MorphoGraphX 2.0 (Strauss et al., 2022).

### Whole plant imaging

Plants were magnified at 50x on a Zeiss Stemi 2000 stereomicroscope. Images were taken with an iPhone Max XS.

### Image quality control

Over the course of live imaging experiments, each day images were inspected for quality and samples were ranked to proceed over many days based on the imaging coverage and signal level. To speed up this process, scripts in ImageJ and MorphoGraphX were implemented. In ImageJ, batch_tiff.py (Supplemental Code 1; https://github.com/kateharline/roeder_lab_projects/tree/master/mgx_scripts) was run on the topmost directory of the imaging files to recursively convert .lsm files from the microscope to .tiff files. Sometimes, ImageJ did not save the z-step in a format that could be read by MorphoGraphX. In this case, multi_resize.py was run in MorphoGraphX to iteratively set the z-step unit across stacks (Video 5; supplemental code 1; https://github.com/kateharline/roeder_lab_projects/tree/master/mgx_scripts). Stacks were then visually inspected in MorphoGraphX for quality. Each stack was examined in the z-direction to ensure a round glow was seen on top indicating the entire top of the sample was captured. For larger samples, each tile was aligned and assembled manually in MorphoGraphX to ensure the entire sample was capitured amongst the individual images. Note, rough assemblies were used for image quality checking. Assembly was repeated more carefully for final image processing.

### Image processing

Most images could be processed on an iMac Pro with Intel Xeon W 3.2 GHz 8 core CPU, 64 GB RAM, Radeon Pro Vega 64 16 GB GPU running Windows 10 through Bootcamp or a VMWare Fusion Linux Virtual Machine running Ubuntu 20.04.3. The largest samples required a PC with AMD Ryzen 9 5950X 3.4 GHz 16 core CPU, 128 GB RAM, EVGA GeForce GTX Titan X 12 GB Superclocked GPU, Ubuntu 20.04.4 LTS. We found that 128 GB of RAM was necessary for processing the large samples, ~7 DAS leaves. The task list of MorphoGraphX processes and respective parameters used to create 2.5D representations of the confocal stacks are enclosed as Supplemental Information. An annotated description of tasks is also enclosed to complement MorphoGraphX documentation for new users. Briefly the image processing steps proceeded as follows. For samples exceeding a single 20X window, tiles were manually aligned and merged in MorphoGraphX. The clipping plane tools were used to visualize and align the stacks in three dimensions. The pixel editor tool was used to erase overlapping regions to a very small sliver at the junction. Then stacks were combined using the merge process. Masks of the confocal stacks were created through 1-3 rounds of Gaussian blurring, then edge detection and closing holes in older samples where masks showed gaps. From these masks, surfaces were created, then the surface that did not contain signal was manually selected and deleted. The confocal signal was then projected onto the surface. Meshes were subdivided once, then subject to 2-3 rounds of auto-segmentation, adaptive mesh subdivision at the new cell borders and projection of the confocal signal back onto the refined mesh. Cell segmentations were manually corrected immediately after segmentation or through the process of manual cell lineage tracing and cell junction correction using the check correspondence process. Meshes from consecutive time points were manually overlaid and cell parents annotated either manually (Video 1-2) or using the semi-automatic parent labeling protocol. Parent tracking quality was assessed using the check correspondence function. Once meshes passed these quality control steps, iterative_growth_and_measures.py (Supplemental Code 1; https://github.com/kateharline/roeder_lab_projects/tree/master/mgx_scripts) was run to calculate growth and cellular parameters and produce heat map representations of the data with standardized parameters across time point comparisons and replicates (Video 4).

### Data analysis

All data processing, analysis and plotting was performed in RStudio (2020; 2021). Scripts used to process the data and create figures are enclosed as Supplemental Information and available at https://github.com/kateharline/live_img_paper, https://github.com/kateharline/roeder_lab_projects/tree/master/imagej_scripts and https://github.com/kateharline/jawd-paper.

### Data availability

Imaging data will be deposited XXXX.

## Funding

Kate Harline was supported by NSF Graduate Research Fellowship (DGE-1650441). This work was funded by NSF MCB-2203275 (AHKR), The Schwartz Research Fund Award (AHKR), and the National Institute Of General Medical Sciences of the National Institutes of Health under Award Number R01GM134037 (AHKR). The content is solely the responsibility of the authors and does not necessarily represent the official views of the National Institutes of Health and other funders.

## Author contributions

K.H. collected, processed and analyzed data. K.H and A.H.K.R. designed experiments, wrote and edited the manuscript.

## Acknowledgements

The authors would like to thank the members of the Roeder and Smith labs for useful discussions and assistance with MorphoGraphX.

---

**Video 1. Parent tracking in MorphoGraphX** Two meshes from the same replicate imaging series are shown in MorphoGraphX. The cell labels are on for the first and second time point’s mesh. The mesh border color is changed for the previous time point, to make cell outlines easier to see. The previous time point is scaled and laid over the subsequent time point. The parent tracking tool ‘Grab label from other surface’ is selected. Previous time point cells are clicked through to transfer parent labels to subsequent time point mesh. White scale bar = 200um (before scaling of the first mesh).

**Video 2. Results of parent tracking in MorphoGraphX** Two meshes from the same replicate imaging series are shown in MorphoGraphX. The cell labels are turned on for the first and second time point’s mesh. The previous time point is scaled and laid over the subsequent time point. The corresponding parent labels are turned on for the second time point’s mesh to demonstrate how the cell labels were transferred onto the successive time point. White scale bar = 200um (before scaling of the first mesh).

**Video 3. Perfluorodecalin mounting solution improves sample vitality over water-based solutions** Animation of undissected and water submersed or dissected and water submersed or dissected and perfluorodecalin submersed samples (left to right). The growth of the first two samples begins to slow and eventually stalls from 5-7 DAS. The perfluorodecalin sample continues to grow. Maximum intensity projections of confocal stacks are shown false colored in green or gray. Red scale bar = 100um.

**Video 4. MorphoGraphX script to rapidly quantify mesh characteristics and capture screenshots** Script parameters are edited to denote which meshes will be run, what measures will be applied, what heatmap images and mesh attributes will be saved. Both intra- and inter-mesh measures can be computed. All measures or only some can be saved as attributes for downstream analysis. The script opens each mesh in time order then conducts measures. The script exits and prompts the user to select cells from which to measure Medial-Lateral distance. Later, the script will exit to prompt the user to arrange the meshes before creating snapshots. The script creates new folders to save attribute maps of measures, snapshots and saves updated meshes.

**Video 5. MorphographX script to resize confocal stack voxels** Confocal stacks exported from ImageJ with z-step incorrectly recorded can appear flat in MorphographX software. Users can designate which files to resize with script parameters. Users can specify voxel dimensions in script parameters. Running multi_resize.py resizes all stacks to voxel dimensions specified by the user in the script file and saves the resized stacks.

## Notes

https://drive.google.com/drive/folders/1zeEszKmO6bKzMwSC1g7yceyOs3kB8eYM?usp=share_link

### Competing Interest Statement

The authors have declared no competing interest.

### Summary of Updates

Add citation for paper with more detailed analysis

