## Supplementary figures and images for "“An optimized pipeline for live imaging whole Arabidopsis leaves at cellular resolution”"

### Video 3

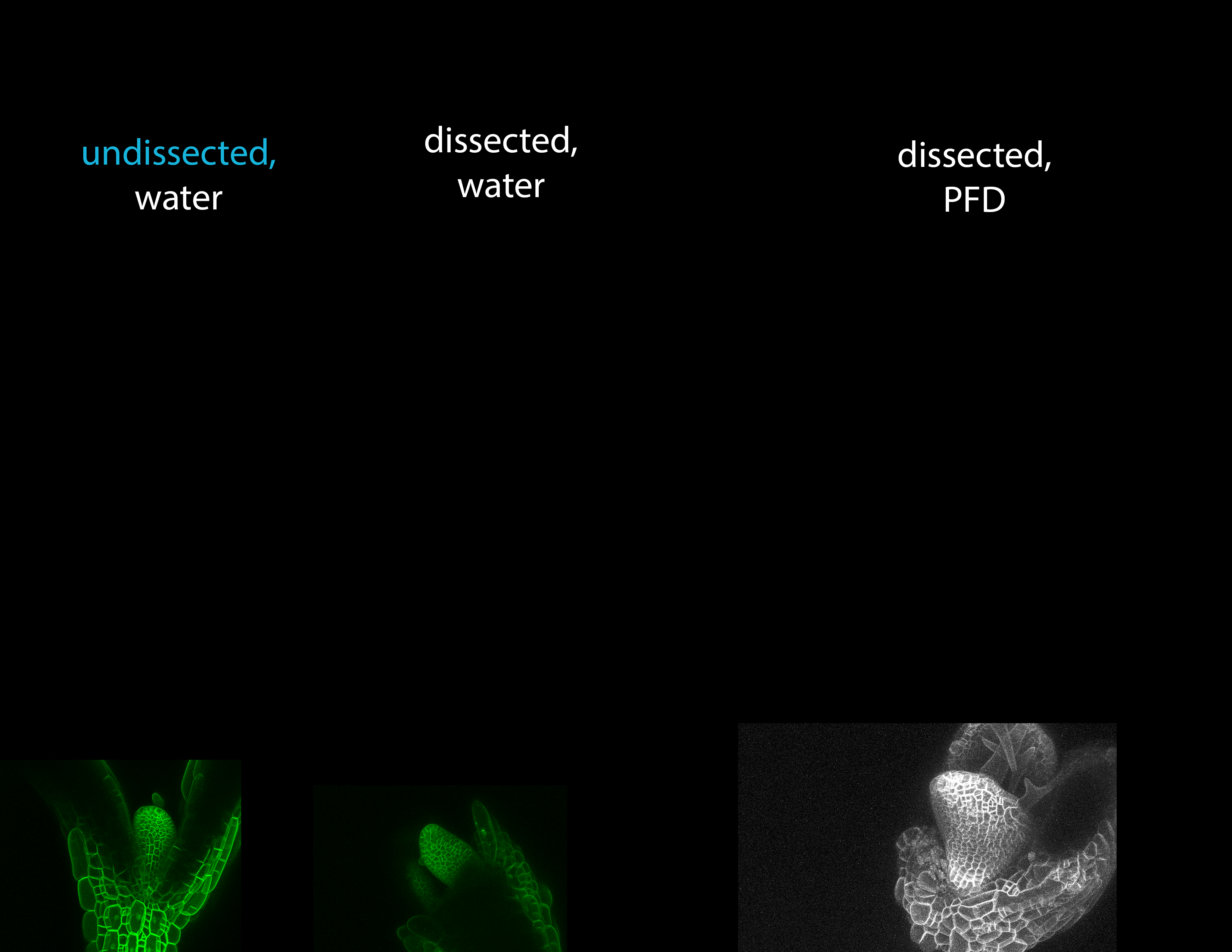
